# Verifying LLM-extracted text with token alignment

**DOI:** 10.64898/2026.02.06.704502

**Authors:** A. Sina Booeshaghi, Aaron Streets

**Affiliations:** Department of Bioengineering, University of California Berkeley, Berkeley, USA; Center for Computational Biology, University of California, Berkeley, USA; Biophysics Graduate Group, University of California, Berkeley, USA; Chan Zuckerberg Biohub – San Francisco, USA

## Abstract

Large language models excel at text extraction, but they sometimes hallucinate. A simple way to avoid hallucinations is to remove any extracted text that does not appear in the original source. This is easy when the extracted text is contiguous (findable with exact string matching), but much harder when it is discontiguous. Techniques for finding discontiguous phrases depend heavily on how the text is split—i.e., how it is tokenized. In this study, we show that splitting text along subword boundaries, with LLM-specific tokenizers, and aligning extracted text with ordered alignment algorithms, improves alignment by about 50% compared to word-level tokenization. To demonstrate this, we introduce the Berkeley Ordered Alignment of Text (`BOAT`) dataset, a modification of the Stanford Question Answering Dataset (`SQuAD`) that includes non-contiguous phrases, and `BIO-BOAT` a biomedical variant built from 51 bioRxiv preprints. We show that text-alignment methods form a partially ordered set, and that ordered alignment is the most practical choice for verifying LLM-extracted text. We implement this approach in `taln`, which enumerates ordinal subword alignments.

## 1 Introduction

Text extraction is a basic task for large language models (LLMs). A user asks the model to extract information from a passage, and the model is expected to return text that actually appears in it. Reliable text extraction is enabling. It allows users from many domains to turn unstructured text—such as a scientific article, historical newspaper, health record, or financial report—into structured data [Gupta et al., 2022, Dunn et al., 2022, Agrawal et al., 2022, Kartchner et al., 2023, Huang et al., 2023, Dagdelen et al., 2024, Zhang et al., 2024, Polak and Morgan, 2024, Chataut et al., 2024, Perot et al., 2024, Dagli et al., 2024, Le Guellec et al., 2024, Liu et al., 2024, Castro et al., 2024, Gartlehner et al., 2024, Pai et al., 2024, Ntinopoulos et al., 2025, Tang et al., 2024, Geevarghese et al., 2024, Schilling-Wilhelmi et al., 2025, Azar et al., 2025, Schwitter, 2025, Qin et al., 2025, Yamagishi et al., 2025, Srivastava et al., 2025, Jiang et al., 2025, Fei et al., 2025, Zhou et al., 2025].

In medicine and biology, accurate text extraction is especially important. For instance, organ procurement organizations rely on lab values and vital signs—^1^often buried inside unstructured clinical notes—to judge whether a potential donor’s organs are suitable [Adam et al., 2024]. Errors introduced during extraction can potentially misinform clinical decisions, and ultimately harm patients.

Fortunately, text extraction can be verified: we can align the extracted text back to its source (Figure 1). For uninterrupted (“contiguous”) phrases, verification is straightforward. An exact match confirms its existence. For non-contiguous phrases, verification is harder; alignment must handle intervening elements. Parenthetical notes, superscripts, symbols, qualifiers, or punctuation may interrupt a logically unified (but practically discontinuous) phrase. For example, the phrase “`natural killer cells`” in Figure 1 cannot be contiguously aligned because the intervening parenthetical expression “`(NK)`” breaks it. Even though the phrase exists, it cannot be found by naive alignment. This situation is particularly acute in biology and medicine where non-contiguous phrases and non-standard nomenclature are highly abundant [Lever et al., 2020, Shinohara et al., 2024].

**Figure 1:**
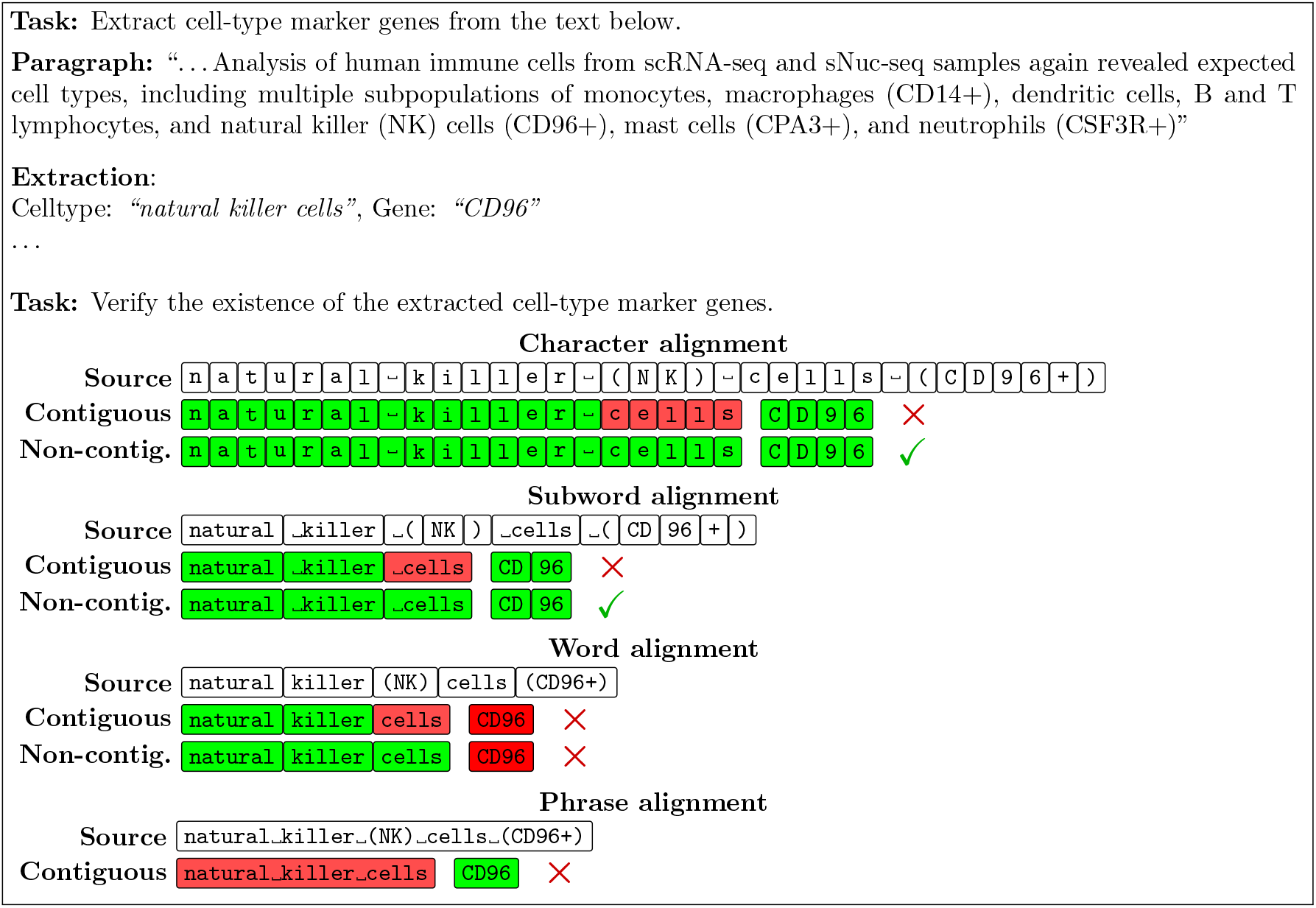
An example of common text-extraction and verification tasks using text from [Hildreth et al., 2021]. LLMs correctly extract marker genes from a sentence, but classical verification tools—that align phrases or words—fail (red). The example cell-type and marker gene are found (green) only when aligned with non-contiguous alignment under subword or character tokenization. Compared to character tokenization, subword tokenization produces fewer tokens resulting in a smaller memory usage.

One way to verify the existence of non-contiguous phrases is with LLM-as-a-judge [Gu et al., 2025]. In this setting, language models are given a passage and extracted piece of text and are prompted to determine its existence. While sometimes effective, LLM-as-a-judge is mismatched to the task. Text is a concrete feature of a passage whereas LLM judgments are semantic and probabilistic distributions over a vocabulary (that includes text from the passage, as well as other text). Therefore, LLMs may approve paraphrases, or hallucinate, making them unsuitable for verifying the existence of text.

Practical issues with LLMs compound this problem. LLM inference is slow, expensive, and non-deterministic.

For example, using GPT5.1 ($1.25 per million input to-kens, $10 per million output tokens pricing as of late 2025), verifying a single 10-character string in a 1,000-character passage costs approximately $0.001 and takes 1-2 seconds. Across 100,000 verifications, this amounts to tens to hundreds of dollars and tens of hours of compute—resources spent replicating what deterministic string matching could achieve instantly at negligible cost.

But as we saw, naive methods are insufficient. More complex methods—that split the phrase and align parts of it—offer an improvement. By splitting the text into units (also known as tokens) and aligning each independently, rather than matching consecutively occurring characters, these methods can align non-contiguous text. Because tokens only need to appear in the same order, non-contiguous matches become possible. Removing the contiguity constraint allows alignment to skip irrelevant parts of a sentence.

Tokens can, in principle, be any unit of text—from individual characters to full words or even multi-word expressions—but token size introduces a fundamental tradeoff [Whittington et al., 2024]. Small tokens (e.g. characters) offer maximal flexibility. Every character can be aligned independently, allowing methods to recover highly fragmented phrases (Figure 1). But this flexibility is computationally expensive. If a phrase has *k* characters and the source has *n* characters (all the same), aligning at the character level can produce up to 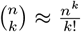 possible non-contiguous matches. For example, aligning a 10-character target (e.g. AAAAAAAAAA to a 1,000 character (`A…A`) source produces roughly 10^23^ possible alignments, compared to only *n − k* + 1 = 991 matches for simple contiguous search. Conversely, very large tokens (e.g. whole sentences) reduce the search space by merging punctuation, symbols, acronyms, or alphanumeric identifiers in ways that may hide LLM-extracted phrases. Thus, token size directly determines what alignments are findable and computationally tractable.

A common tokenization method is word-level: text is split on the spaces between characters and tokens are whole words. But word-level tokenization breaks down in scientific text. As shown in Figure 1, if an LLM extracts the gene name “`CD96`” from the phrase “`natural killer (NK) cells (CD96+)`”, word-level alignment fails because that tokenization merges the gene name with parentheses, and a plus sign. This situation is quite common. In biological text, gene names often contain characters like parentheses, plus signs, and mixed alphanumeric formatting that are inconsistently attached to genes, causing word-level methods to miss alignments.

In this work, we study how tokenization and alignment strategy affect the recovery of missing alignments resulting from non-contiguous phrases. We introduce two benchmark datasets, formalize text alignment to motivate a practical alignment method, and evaluate the impact of tokenization with this method. Based on these benchmarks, we release an open-source tool, called `taln`, for accurate non-contiguous text alignment.

## 2 Results

We constructed two benchmark datasets: `BOAT`, derived from `SQuAD` [Rajpurkar et al., 2016, 2018], and `BIOBOAT`, a biomedical extension built from 51 bioRxiv preprints (Table 1). Using these datasets, and a formalization of alignment methods, we show that ordered alignment is most practical for aligning non-contiguous phrases and that tokenization greatly impacts alignment accuracy. Furthermore, subword tokenization [Sennrich et al., 2016] recovers substantially more alignments than word-level tokenization while remaining computationally practical.

**Table 1:** The Berkeley Ordered Alignment of Text (BOAT) dataset. consists of multiple human-annotated text extractions derived from the `SQuAD` dataset. Each example includes a target text and a corresponding source text, where the target can be identified by direct string matching in the source. `BOAT` has two parts: a contiguous one, where the full target appears in the source, and a non-contiguous one, where part of the target is ablated and the target cannot be found contiguously. We report the average character (Char), whitespace-delimited word (WS), and subword token (Tok) lengths of both source and target texts. WS corresponds to word-level, and Tok to subword tokenization. Note that a small fraction of targets that are present in the source cannot be found with non-contiguous subword tokenization due to incorrectly set tokenization boundaries present in the `SQuAD` dataset. We also report the same statistics for the `BIO-BOAT` dataset.

| BOAT | N | N<sub>0-align</sub> (%) | $\mu_{\text{align}}$ | Text | Chars | WS | Tok |
| --- | --- | --- | --- | --- | --- | --- | --- |
| Contiguous | 35,080 | 159 (0.4) | 8 | source | 764 | 121 | 158 |
|  |  |  |  | target | 38 | 6 | 8 |
| Non-contiguous | 35,080 | 198 (0.6) | 7 | source | 764 | 121 | 158 |
|  |  |  |  | target | 33 | 5 | 7 |
| Contiguous (BIO) | 5,316 | 171 (3.2) | 94 | source | 607 | 91 | 132 |
|  |  |  |  | target | 97 | 14 | 20 |
| Non-contiguous (BIO) | 5,316 | 584 (11.0) | 89 | source | 607 | 91 | 132 |
|  |  |  |  | target | 90 | 13 | 18 |

### 2.1 Dataset design

`BOAT` consists of 35,080 source–target pairs derived from `SQuAD`, where each target aligns contiguously in its source passage. Sources average 764 characters (121 whitespace tokens; 158 subword tokens), and targets average 38 characters (Table 1). To evaluate non-contiguous alignment, we constructed a variant in which one interior token is removed from each target, ensuring that the resulting phrase can no longer be found by exact contiguous alignment.

To test whether alignment behavior generalizes beyond general-domain text, we constructed `BIO-BOAT`, containing 5,316 source–target pairs extracted from 51 bioRxiv preprints, with both contiguous and non-contiguous variants (Methods) [van Vliet et al., 2022, Cobo-López et al., 2023, Meyer et al., 2023, Zimyanin et al., 2023, Singer et al., 2024, Shen et al., 2024, Frolov et al., 2024, Ryan et al., 2024, Pfalzgraf et al., 2024, Onishi et al., 2024, Wondim et al., 2024, Crnkovic et al., 2024, Kallenborn et al., 2024, Peroutka-Bigus et al., 2024, Azim et al., 2024, Bhalla et al., 2024, Jones et al., 2024, Fei et al., 2024, Combredet and Brunet, 2024, Simmonds et al., 2025, Adhinarta et al., 2025, Elangovan et al., 2025, Tang et al., 2025, Maya-Miles et al., 2025, Samame-Caramutti et al., 2025, Nguyen et al., 2025, Terstege and Epp, 2025, Chen et al., 2025, Li et al., 2025, Kattner et al., 2025, Shruti and Vijay Prakash, 2025, Velázquez et al., 2025, Rull, 2025, Dölker et al., 2025, Keikhosravi et al., 2025, Agazzi et al., 2025, Choltus et al., 2025, Maestro et al., 2025, Widney et al., 2025, Smith et al., 2025, Marín, 2025, Dadonaite et al., 2025, Green et al., 2025, Sülzen et al., 2025, Pearce et al., 2025, Biller et al., 2025, Laub et al., 2025, Johnson et al., 2025, Na et al., 2025, Tozzi, 2025]. Biomedical text introduces denser punctuation, symbols, and alphanumeric identifiers, making it a stringent test for alignment methods and tokenizers (which are often built from general-domain texts.)

### 2.2 Alignment algorithms

Many alignment methods exist for verifying extracted text, from exact string matching to algorithms that allow gaps or reorderings. To compare them, we formalized alignment as a map from target token positions to source token positions (Supplementary Note).

Under this formalization, alignment methods form a hierarchy defined by the constraints they impose on this map. Contiguous alignment requires adjacent tokens to match. Ordered alignment drops adjacency but retains order. Permutation alignment drops order but requires a one-to-one correspondence. General rearrangement imposes no constraints. Each relaxation admits more valid alignments, but the number of possible alignments grows rapidly, from linear (contiguous) to factorial (permutation) to exponential (rearrangement).

Ordered alignment sits at a practical point in this hierarchy: it recovers non-contiguous phrases while keeping the number of alignments manageable. In practice, relaxing constraints beyond ordered alignment produces spurious alignments, making permutation and rearrangement impractical. We therefore focus our study on ordered alignment and implement it in a tool called `taln`, short for **T**oken **Al**igner. `taln` is a Python library and command-line tool that returns all order-preserving alignments between a target and source. `taln` is tokenizer-agnostic and supports various tokenization strategies.

### 2.3 Text Tokenization

Having selected an alignment algorithm, we next asked how tokenization affects alignment accuracy. We compared naive phrase matching, word-level tokenization (whitespace), and subword tokenization using the `cl100k base` tokenizer (Methods). We benchmarked `taln`, longest common subsequence (LCS), and Python’s `difflib.SequenceMatcher` on both contiguous and non-contiguous variants of `BOAT` and `BIO-BOAT` (Figure 2).

**Figure 2:**
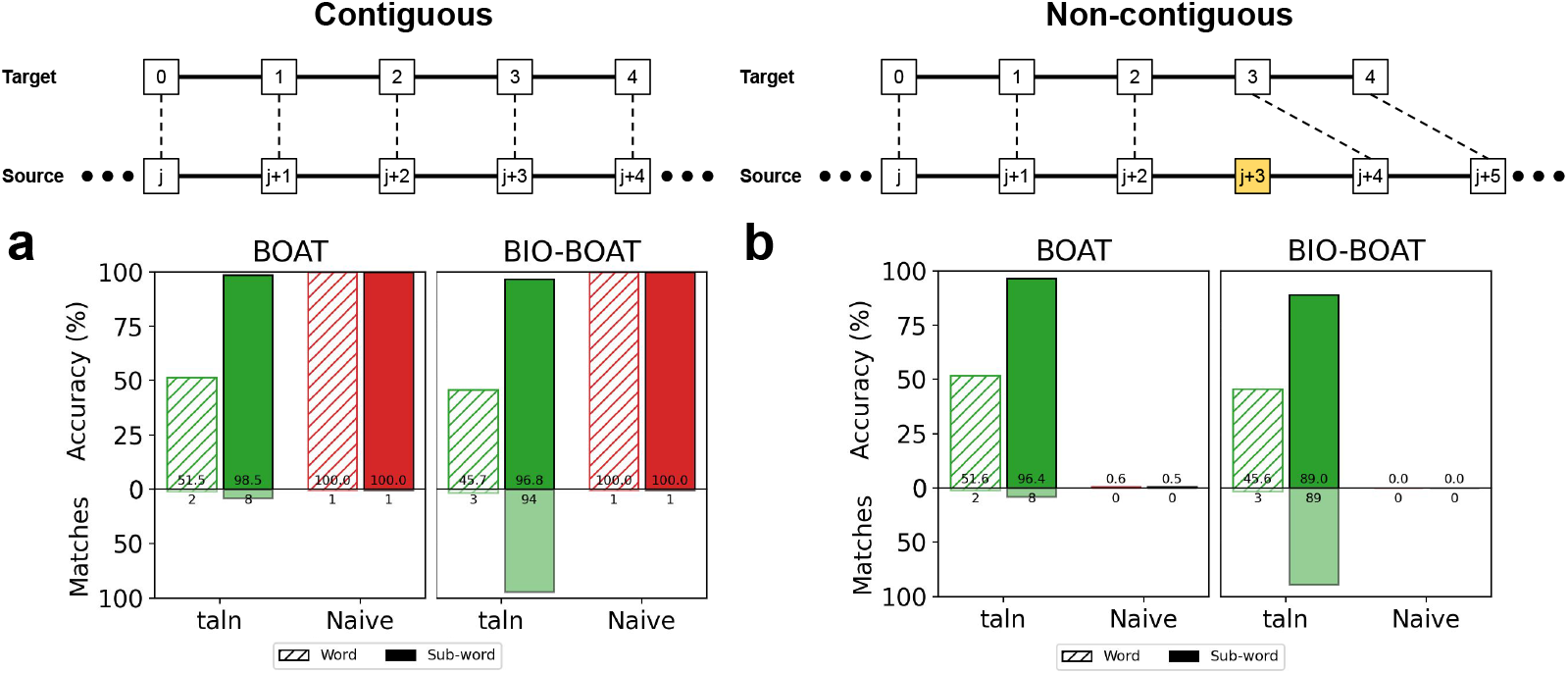
Contiguous and non-contiguous alignment. **a**. Contiguous alignment accuracy for all samples in the `BOAT` (left) and `BIO-BOAT` (right) datasets for `taln` (green) vs. Naive alignment (red) tokenized at the word (striped) and subword (solid) level as well as the average number of alignment matches for each method (below). **b**. Non-contiguous alignment accuracy and enumeration for `taln` and Naive alignment on the non-contiguous `BOAT` (left) and `BIO-BOAT` (right) datasets.

#### 2.3.1 Contiguous text

On contiguous phrases from `BOAT`, naive alignment recovered all targets, as expected. In contrast, under word-level tokenization, nearly half of targets were missed by all ordered-alignment methods despite being fully contiguous (47–58%, depending on the tool; Figure 2a and Supplementary Figure S1a, striped).

These failures arose from inconsistencies introduced by whitespace tokenization, where punctuation or quotation marks are inconsistently attached to neighboring words, preventing correct token-level alignment. For example, punctuation like quotation marks and commas are often omitted from the beginning or end of `SQuAD` answers (e.g. `a lawless place’’ or ‘‘a place under no jurisdiction`) even when present in the passage (`…of ‘‘a lawless place’’ or ‘‘a place under no jurisdiction’’…`). When split by whitespace, the missing adjacent quotes prevent correct alignment, causing otherwise contiguous phrases to be missed.

To address these missing alignments, we repeated the evaluation using subword tokenization (Methods). Subword tokenization largely resolves this issue. By splitting words into smaller fragments, subword tokenization separates punctuation and symbols from meaningful text—e.g., (`of ‘‘a lawless`) splits into (`of, ‘‘, a, law, less`). As a result, alignment accuracy increased to 98–99% across all methods (Figure 2a and Supplementary Figure S1a, solid), demonstrating that tokenization—not the alignment algorithm alone—also contributes to alignment accuracy.

#### 2.3.2 Non-contiguous text

To evaluate alignment performance on non-contiguous phrases, we used the non-contiguous `BOAT` dataset. In this setting, naive alignment failed almost entirely (0.6%). Under word-level tokenization, ordered alignment recovered only about half of targets (51–52%; Figure 2b and Supplementary Figure S1c, striped).

Failures arose from the same issue observed in the contiguous setting: punctuation, symbols, or alphanumeric patterns are inconsistently segmented at the word level, making ordered alignment difficult once phrases are tokenized.

Subword tokenization dramatically improved performance. With subword tokens, ordered alignment recovered over 96% of non-contiguous targets across all tools, including `taln`, LCS, and `difflib` (Figure 2b and Supplementary Figure S1c, solid). This shows that allowing gaps is necessary but not sufficient; alignment of non-contiguous phrases depends on the granularity of subwords defined by the tokenization strategy.

#### 2.3.3 Generalization to biomedical text

To test whether these results generalize to biomedical text, we repeated the evaluation on `BIO-BOAT` (Figure 2a,b, right; Supplementary Figure S1b,d).

The same alignment patterns hold. For contiguous phrases, word-level tokenization missed roughly half of targets, while subword tokenization recovered over 97%. For non-contiguous phrases, word-level tokenization recovered only 45–46% of targets, whereas subword tokenization improved accuracy to approximately 89% (Figure 2).

The slightly reduced alignment on `BIO-BOAT` likely reflects the complexity of biomedical vocabulary, where gene names and chemical identifiers are often split inconsistently and therefore cannot be reliably captured by a tokenizer trained on general-domain text.

### 2.4 Alignment enumeration

Unlike LCS and `difflib`, `taln` enumerates all valid alignments rather than returning a single match. This matters when extracted phrases appear multiple times in a passage—common in scientific text where repeated entities (gene names, conditions, acronyms) may indicate importance [Swayamdipta et al., 2018, Zhang et al., 2023, Moon et al., 2023, Wang et al., 2025].

On `BOAT`, `taln` returned an average of 8 alignments per contiguous target and 7 per non-contiguous target (Table 1), compared to a single alignment produced by standard tools.

On `BIO-BOAT`, `taln` returned an average of 95 alignments per contiguous target and 89 per non-contiguous target. This increase likley reflects finer to-ken resolution caused by a mismatch between biomedical text and general-domain tokenizers. For example, phrases such as “`secrete the anti-angiogenic isoforms VEGF-A165b`” are split into multiple tokens; the gene “`VEGF-A165b`” alone is decomposed into five tokens, which combinatorially increases the number of potential alignments when using `taln`.

### 2.5 Runtime & Memory

Finally, we evaluated runtime and worst-case memory requirements (Figure 3). While naive alignment is fastest, all ordered alignment methods operate in the millisecond range for typical passage lengths (*∼* 1,000 tokens). `taln`, LCS, and `difflib` scale approximately linearly with source length, remaining practical for standard short-passage alignments (Supplementary Figure S3).

**Figure 3:**
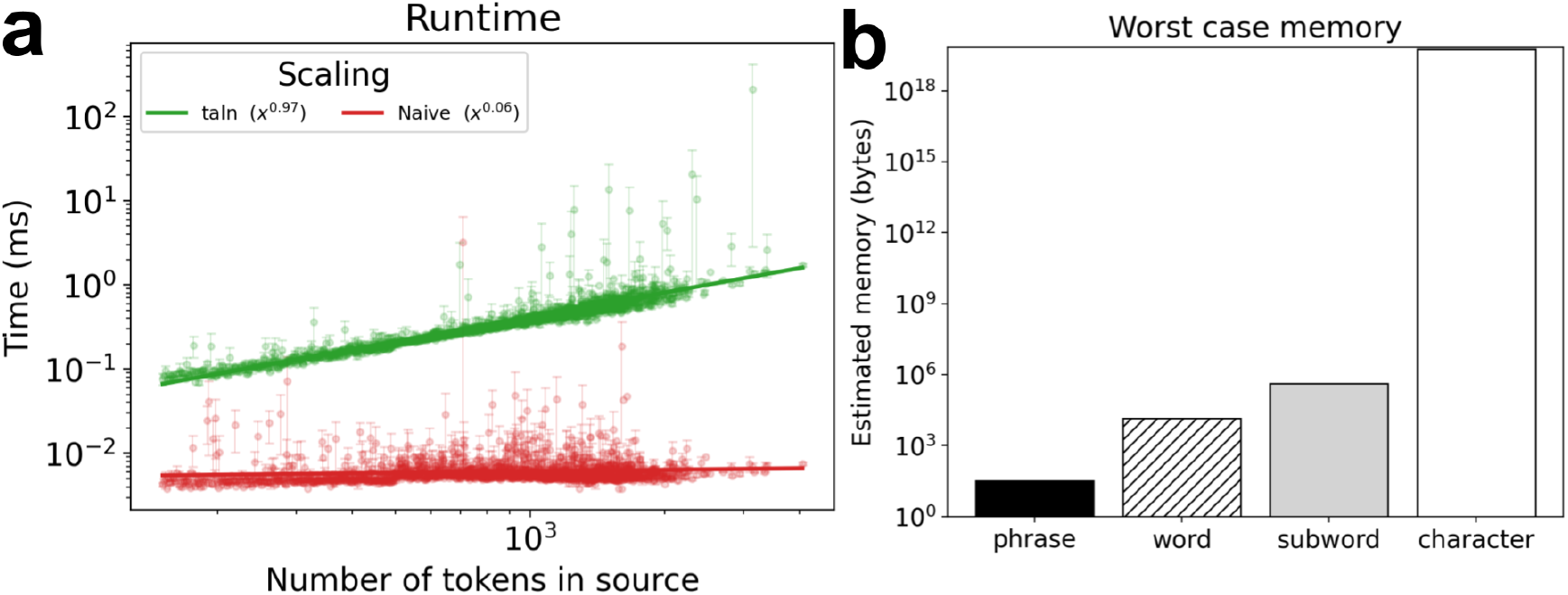
Runtime & Memory: **a**. Runtime of `taln` vs. naive phrase alignment as a function of source length. The inset power law shows the runtime scaling as a function of the number of tokens in the source. **b**. Estimated worst-case memory requirements for phrase (black), word (striped), subword (gray), and character (white) tokenization.

Memory usage, however, varies sharply with the granularity of tokenization. Character-level tokenization incurs orders-of-magnitude higher memory costs, making it impractical despite its flexibility. Subword tokenization offers a practical compromise, retaining most of the alignment flexibility of character-level methods while remaining computationally feasible.

## 3 Discussion

LLMs are powerful tools for data mining. They can ingest massive amounts of text and rapidly synthesize, extract, and return specific pieces of information. But unlike other data-mining tasks, such as summarization, text extraction has a strict constraint on the generated text: the output must already exist in the input. Humans can often verify that this constraint is not violated, but doing so at scale becomes prohibitive. This is where text-alignment algorithms can assist human evaluators by providing automated, verifiable checks. Our results show that this verification constraint can be satisfied deterministically using token-based alignment, without requiring semantic judgment by an LLM.

Alignment algorithms are well suited to this verification task, but traditional text alignment algorithms and tokenization strategies struggle with scientific text— despite their long history and development in disparate fields [Hirschberg, 1975, Knuth et al., 1977, Melsted et al., 2021, Sennrich et al., 2016, Sullivan et al., 2024]. Extracted phrases are often interrupted by symbols, qualifiers, or parenthetical statements. Exact phrase matching therefore misses many valid alignments. Alignment methods that allow gaps or skips between aligned segments offer a practical solution, but they introduce a new question: what constitutes a meaningful “part” of a phrase?

This work shows that subword tokenization—used by many modern LLMs for training and inference—provides a natural answer. When combined with ordered alignment methods such as `taln`, it enables reliable alignment even when their contiguity is unknown. Subword tokenization improves accuracy in both contiguous and non-contiguous settings, and `taln`, by enumerating all valid alignments, recovers matches that standard tools miss, while remaining fast and deterministic compared to LLM-based verification approaches.

However, enumerating many valid alignments introduces a new challenge: selecting the most meaningful one. As recall improves, specificity becomes the limiting factor. Techniques for selecting among numerous valid alignments are a promising direction to improve the specificity of `taln`, for example using approaches based on token embedding radius (or related selection criteria, Supplementary Note.) While not discussed here, methods for verifying the existence of LLM-extracted phrases with semantic (instead of exact) matching are also a promising direction.

The selection challenge is amplified by domain-specific differences. The lower accuracy, and greater alignments, on `BIO-BOAT` compared to `BOAT` likely reflects more complex tokenization boundaries in biomedical text—gene names, chemical formulas, and mixed alphanumeric identifiers are common and often split inconsistently even by subword tokenizers, suggesting the need for domain-specific tokenizers to further improve alignment specificity.

Together, these results demonstrate that alignment algorithms and tokenization matter for verifying LLM-extracted text, and how methods from computational biology and linguistics can be combined to develop useful verification tools. Practically speaking, we envision non-contiguous alignment algorithms used with subword tokenization in text-extraction workflows where refined text alignment can be performed: passages can be extracted contiguously from large documents, and relevant— possibly non-contiguous—text can then be aligned to those passages, augmenting human ability to parse documents and ground results at scale.

## A Impact Statement

Large language models are increasingly used to extract structured information from text in scientific research, medicine, and other data-intensive fields. This use is attractive because models can process volumes of text far beyond human capacity. However, extraction is only useful if the returned text actually appears in the source—i.e. it is not hallucinated. In high-stakes settings such as clinical decision support or scientific data curation, failures of verification can propagate errors and undermine trust.

This work addresses verification by improving how extracted text is aligned back to its source. We show that token-based alignment is most effective when it uses subword tokenization, which matches the representation used by LLMs, rather than whitespace- or character-based tokenization. On our benchmarks, this choice recovers about 50% more alignments than word-level while being more memory efficient than character level. By providing deterministic and transparent alignment algorithms, this work supports more trustworthy use of LLM-based text extraction in scientific and clinical workflows, and reduces reliance on probabilistic or fragile verification methods.

## B Methods

### B.1 BOAT Building

We derived `BOAT` from the Stanford Question Answering Dataset (`SQuAD`) v2.0. Each `SQuAD` example provides a passage (source), an answer string (target), and an answer start index (`answer start`) locating the target as a contiguous character sequence in the source.

We first flattened `SQuAD` into records with fields `source`, `target`, and `idx start` (plus metadata such as title and question identifiers). We retained records where naive contiguous string match occurs: `source [idx start:idx start+len(target)]`=`target`. We then deduplicate on `(source, target, idx start)`.

To make character indices consistent with subword tokenization, we applied a “shift” normalization: if the character immediately preceding `idx start` is a space, we prepend a space to the target and decrement `idx start` by one. This ensures that leading whitespace, when present in the source, is part of the target phrase; a feature that is often seen in substring tokenization vocabularies. Note this does not affect whitespace tokenization since the space is prepended to the string.

Because identical `(source, target)` pairs can occur with multiple valid start indices (e.g., repeated mentions in a passage), we grouped by `(source, target)` and stored `idx start` as a set of start positions. In downstream analyses we restricted to multi-word targets to focus on phrases where ablating an interior token introduces a meaningful non-contiguity.

We constructed two `BOAT` variants: Contiguous `BOAT` and Non-contiguous `BOAT`. Contiguous `BOAT` consists of the original `(source, target)` pairs and their `idx start` sets. Non-contiguous `BOAT`was constructed by removing, for each `(source, target)` pair, one interior token of the target to create a non-contiguous variant. In the benchmarking notebook this ablation removes one interior whitespace-delimited word. In the dataset construction notebook we also generate a subword-token ablation (removing one interior subword token).

#### B.1.1 BIO-BOAT**Building**

We downloaded 51 papers from the bioRxiv AWS bucket `s3://biorxiv-src-monthly/Current content/`, dated January 2025 and licensed under CC BY 4.0. We converted the manuscript XML to Markdown and ran `taln extract` to generate source–target pairs, where each target is a contiguous span of the source. We then generated non-contiguous `BIO-BOAT` in the same manner as `BOAT` described above.

### B.2 Text Tokenization

We evaluated alignment under two tokenization schemes: word (whitespace) tokens and subword tokens. Word tokens are maximal non-whitespace runs found by regex matching. Each token retains its character start and end offsets in the source string. Subword tokens are produced by the OpenAI `cl100k base` tokenizer (via `tiktoken`). We decode tokens with offsets to recover character start and end positions for each token.

Before tokenization, source and target text are normalized by applying Unicode normalization and ASCII transliteration (`unidecode`), mapping several common punctuation/symbol variants to ASCII, and collapsing whitespace and newlines. Because token offsets are defined on the normalized string, we exclude examples where normalization changes string length.

### B.3 Alignment Algorithms

Given a source sequence of tokens and a target sequence of tokens, we evaluate both contiguous and non-contiguous alignment methods: naive phrase matching, longest common subsequence (LCS), `difflib`, and `taln`.

Naive matching checks whether `target` occurs as an exact substring of `source` and returns all matches. LCS computes an order-preserving alignment using dynamic programming and returns the matched token positions (whitespace or subword). `difflib` uses Python’s `difflib.SequenceMatcher` over tokens (whitespace or subword) to identify and return matching tokens.

`taln` builds an index of token *n*-grams from the source (with *k* = 1 in this work), aligns each target token to all occurrences of that token in the source, and then enumerates all order-preserving alignments. Each alignment is returned as a list of token objects with character offsets; multiple valid alignments are enumerated explicitly.

When benchmarking alignment, we report two complementary metrics: reconstruction accuracy (Figure S1) and localization accuracy (Figure S2). Reconstruction asks whether a method can recover the target string, independent of where it came from in the source. Concretely, reconstruction succeeds if at least one returned alignment reconstructs the target exactly (after applying the same normalization used for tokenization). Localization asks whether a method recovers the ground-truth target (positionally). For each fully reconstructed alignment, we take its bounding position (`start idx`, `end idx`). Localization succeeds if any reconstruction matches the positions of the target (*s, s* + |`target`|) for some *s ∈* `idx start`.

For non-contiguous alignments, localization accuracy is defined by the bounding character indices: an alignment is correct if the left and right character boundaries (`start idx`, `end idx`) of at least one fully reconstructed alignment match the expected target.

### B.4 Runtime Results

We measure per-sample runtime for each alignment method by wall-clock timing around the alignment call on each example, reporting elapsed time in milliseconds. Runtime and accuracy are summarized as a function of source length (measured in tokens) to illustrate scaling behavior with input size.

## C Data and Code Availability

The `SQuAD` datasets can be found here https://rajpurkar.github.io/SQuAD-explorer/. The `taln` tool and the `BOAT` and `BIO-BOAT` datasets can be found here: https://github.com/sbooeshaghi/taln.

## D Acknowledgments

We thank Lior Pachter and Gennady Gorin for helpful feedback on the presentation of the main text as well as for comments on the Supplementary Note. We thank Nithya Appannagaari for her work—on verifying the existence of extracted marker genes with A.S.B.—that partly inspired the creation of `taln`. We thank Hanna Krasowski, Lauren Malek, and Sewon Min for helpful discussions and Hanna Krasowski for contributing improvements to `taln`. We also thank the Howard Hughes Medical Institute for supporting A.S.B. through the Hanna H. Gray Fellows program.

## E Author Contributions

A.S.B. conceived the project, developed the benchmarks, implemented the `taln` software and analysis code, and wrote the manuscript. A.S. reviewed the manuscript.

## G Supplementary Figures

**Figure S1:**
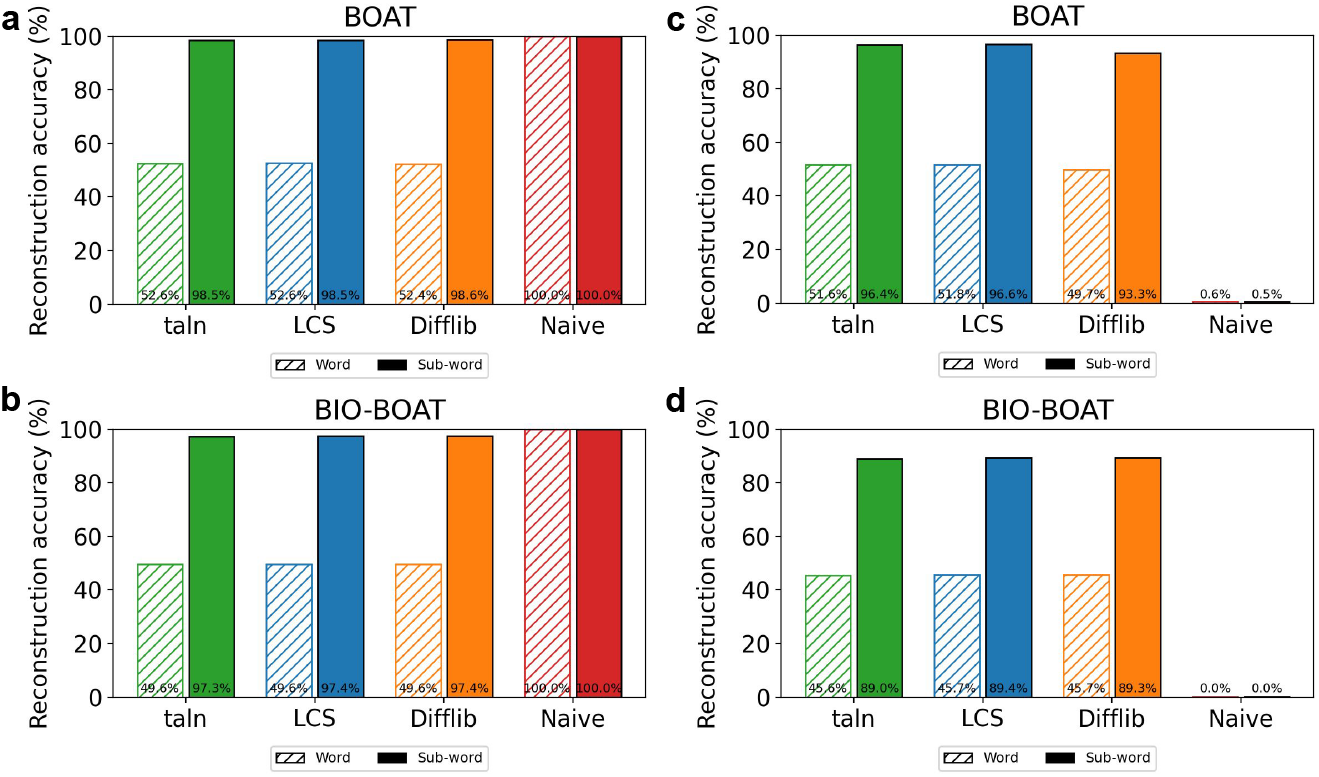
Reconstruction accuracy for contiguous and non-contiguous alignment. Contiguous alignment reconstruction accuracy for all samples in the `BOAT` (a) and `BIO-BOAT` (b) dataset for `taln` (green), LCS (blue), Difflib (orange), and Naive alignment (red) tokenized at the word (striped) and subword (solid). Non-contiguous alignment reconstruction accuracy all tools on the non-contiguous `BOAT` (c) and `BIO-BOAT` dataset (d).

**Figure S2:**
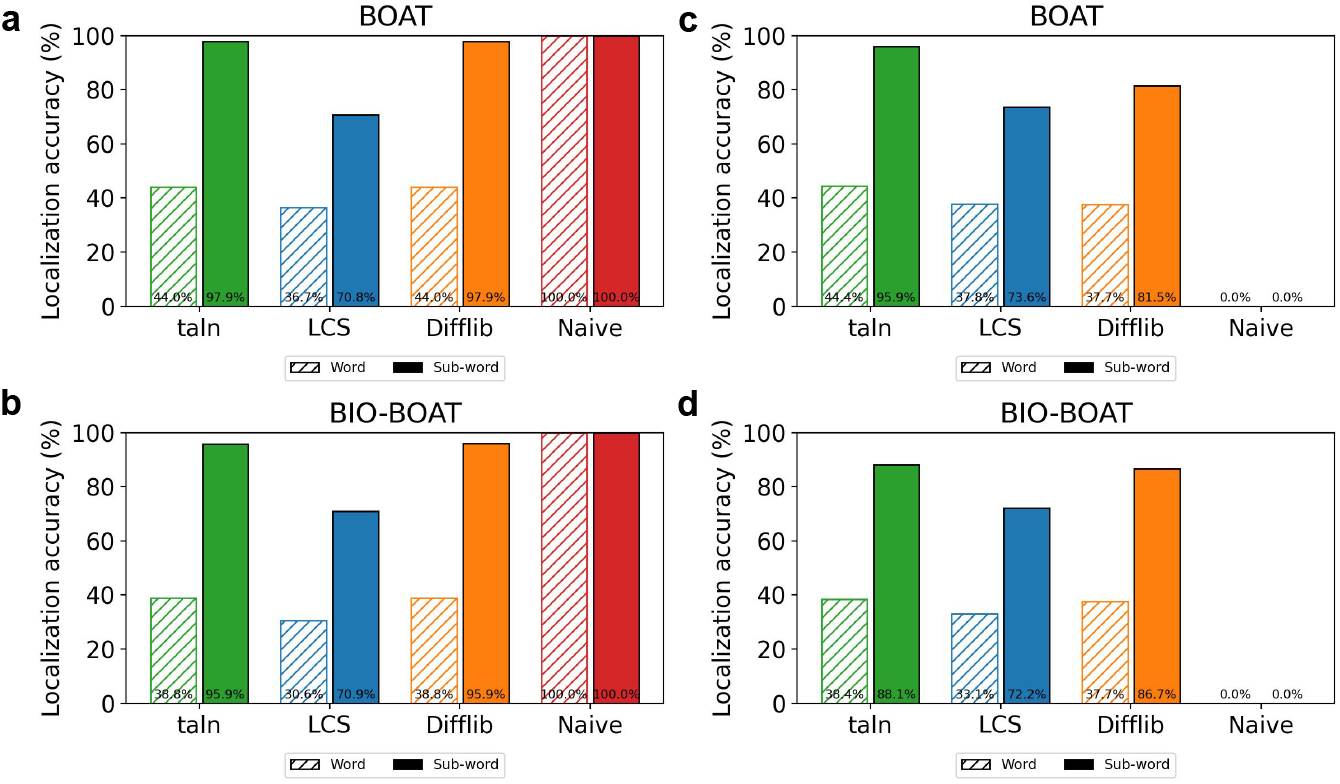
Localization accuracy for contiguous and non-contiguous alignment. Contiguous alignment localization accuracy for all samples in the `BOAT`(a) and `BIO-BOAT` (b) dataset for `taln` (green), LCS (blue), Difflib (orange), and Naive alignment (red) tokenized at the word (striped) and subword (solid). Non-contiguous alignment localization accuracy all tools on the non-contiguous `BOAT` (c) and `BIO-BOAT` dataset (d).

**Figure S3:**
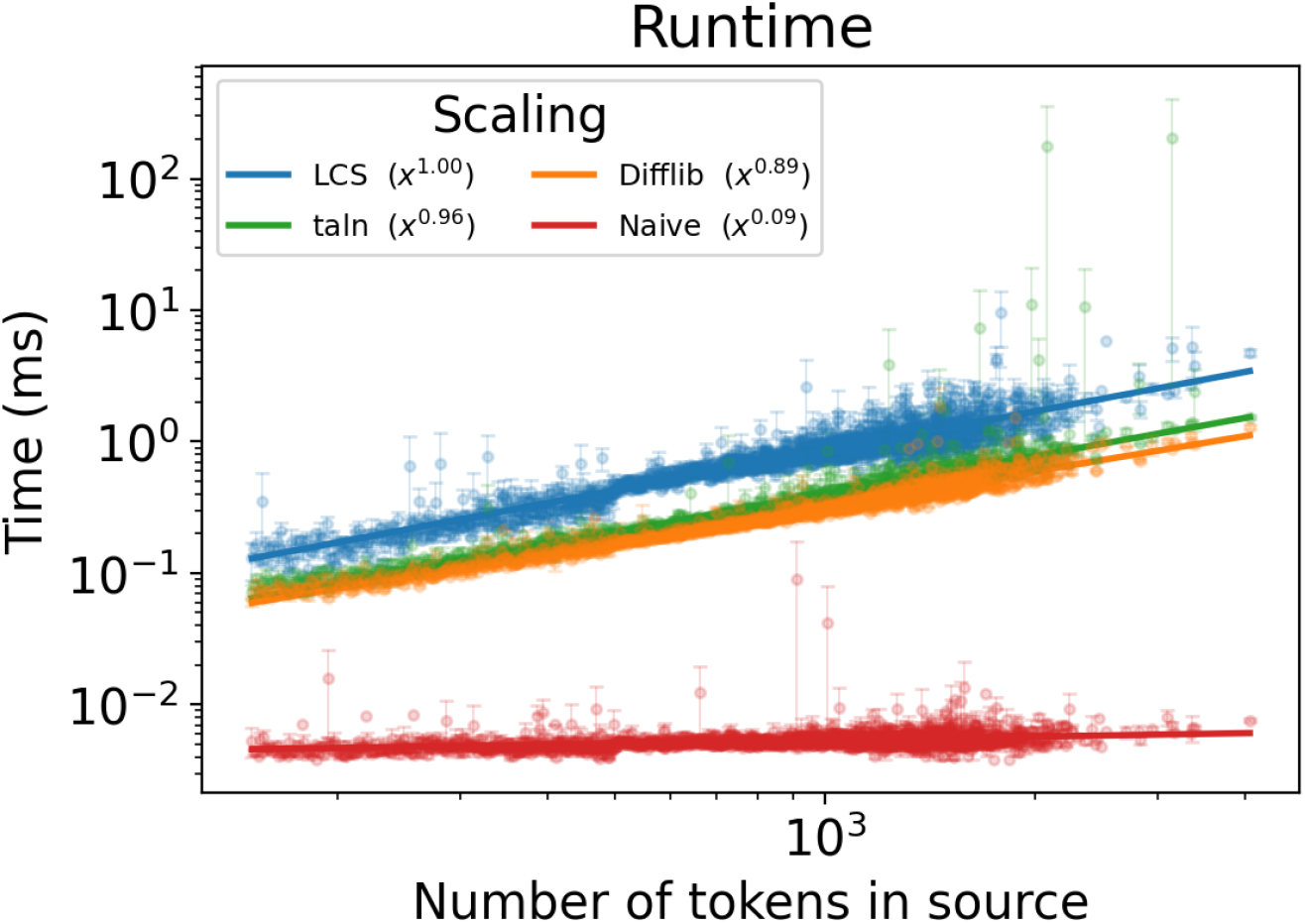
Runtime of various text alignment tools. Log–log plot of runtime (ms) versus number of tokens in the source for four alignment methods. Points show individual source-target pairs; lines show power-law fits with exponents indicated in the legend. LCS and `taln` scale roughly linearly, Difflib is slightly sublinear, and the naive baseline is nearly constant.

## Footnotes

1 The em dashes reflect our stylistic preference and should not be interpreted as evidence of AI-assisted text generation.

