## Supplementary Note for "Verifying LLM-extracted text with token alignment"

Sequence alignment has a long history of algorithm development in computational linguistics and genomics. Classical methods include text matching algorithms like Knuth–Morris–Pratt [Knuth et al., 1977] for exact substring detection, dynamic-programming aligners for gapped and approximate matches [Needleman and Wunsch, 1970, Smith and Waterman, 1981], pseudoalignment methods for transcriptomics [Bray et al., 2016], and genome rearrangement models for studying structural variation [Hannenhalli and Pevzner, 1999, Tesler, 2002]. Although these tools are used in different domains, they share a common goal: relate one sequence to another through alignment.

Here we develop a unified framework, that makes this connection explicit. We treat every alignment problem as the search for a function $f$ mapping positions in one sequence to positions in another under a chosen set of constraints, inspired by the formalism put forth in [S. Schwartz and Pachter, 2007]. Contiguous matches, ordered subsequences, pseudoalignments, and rearrangements all appear as special cases distinguished only by which constraints on $f$ are enforced or relaxed.

This framework reveals a natural hierarchy. By comparing constraint families, we obtain a partially ordered set over alignment classes, ordered from the most restrictive (exact substring matching) to the least (arbitrary realignments). From this structure, ordered alignment naturally falls out as the most permissive alignment strategy, for text, under practical computational constraints (number of alignments). It sits above classical (contiguous) text alignment, and below permutation alignment. In the main text, we realize this alignment strategy, with subword token sequences, to verify the existence of text extracted by large language models.

#### 1 Results

Text alignment is the process of finding a map between two sequences, the target and the source. If such a map exists, then the target is said to fully align to the source—such as `CAT` aligning to `SCATTER`.

This example is intuitive and simple but makes two implicit assumptions. The first is that the alignment is contiguous—that every aligned character must directly follow the previous. The second is that the objects being aligned are characters themselves (instead of other features like groups of characters, $n$-grams).

By formalizing text alignment and making these assumptions explicit, we can more easily compare complex alignment algorithms. The differences between them get reduced to choices of the alignment objects (tokens) and the constraints on the alignment itself. With this formalization we find that contiguous text alignment is a special case of pseudoalignment.

##### 1.1 Ordered alignment

Suppose we have a source sequence of characters $\mathbf{s} = (s_1, \ldots, s_n)$ and a target sequence of characters $\mathbf{t} = (t_1, \ldots, t_m)$. An order-preserving alignment map is a function $f : \{1, \ldots, m\} \to \{1, \ldots, n\}$, which maps each position of the target to a position in the source, subject to an order constraint on $f$.

If we only care that the alignment is ordered (permitting gaps) then the constraint family is the set of functions $f(1) < f(2) < \cdots < f(m)$, $s_{f(j)} = t_j$, for all $j$. If such an $f$ exists, then $\mathbf{t}$ appears in $\mathbf{s}$ as a (possibly non-contiguous) ordered subsequence. The set of all maps is given by the set of functions:

$$\mathcal{E}_{\mathrm{MA}}(\mathbf{t}, \mathbf{s}) = \{f : f(1) < \cdots < f(m),\ s_{f(j)} = t_j\ \forall j\}. \quad (1)$$

**Example: Character alignment.** Consider $s = \texttt{SCATTER}$ and $t = \texttt{CAT}$. The positions of C and A in $s$ are fixed to 2 and 3, while the final T may align to either $s_4$ or $s_5$.

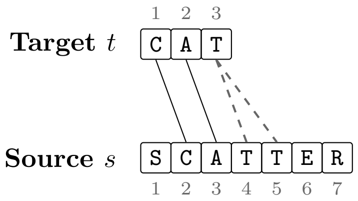


The ambiguity in the final T leads to multiple alignments $\mathcal{E}_{\mathrm{MA}}(\mathbf{t}, \mathbf{s}) = \{f : (f(1), f(2), f(3)) \in \{(2,3,4), (2,3,5)\}\}$, whereas the simple example above (contiguous alignment) permits only one valid map.

#### 1.2 General alignment

With the example above in mind, we now put forth a framework for classifying alignment methods. Let $\Sigma$ be a character alphabet, let $V$ be a vocabulary of tokens, and let $\tau : \Sigma^* \to V^*$ be a tokenizer that maps sequences of characters to sequences of tokens. For a source $S \in \Sigma^*$ and a target $T \in \Sigma^*$, define

$$\tau(S) = \mathbf{s} = (s_1, \ldots, s_n), \tag{2}$$
$$\tau(T) = \mathbf{t} = (t_1, \ldots, t_m). \tag{3}$$

Given a source $\mathbf{s}$ and target $\mathbf{t}$, an alignment map is a function

$$f : \{1, \ldots, m\} \to \{1, \ldots, n\}, \tag{4}$$

which maps each position of the target to a position in the source.

Different alignment tasks impose different requirements on how $f$ may behave. We refer to any collection of such requirements as a constraint family $\mathcal{C}$. Typical constraints include monotonicity, contiguity, distance bounds, membership conditions, and cardinality properties of $f$. A map $f$ is injective if distinct target positions never map to the same source position (no duplication), and surjective if every source position is used by at least one target position (no deletion). A bijective map is both injective and surjective and represents a one-to-one correspondence between source and target positions.

Given a constraint family $\mathcal{C}$, the set of all valid maps of $\mathbf{t}$ into $\mathbf{s}$ is

$$\mathcal{E}_{\mathcal{C}}(\mathbf{t}, \mathbf{s}) = \{f : f \text{ satisfies } \mathcal{C} \text{ and } s_{f(j)} = t_j \ \ \forall j\}, \tag{5}$$

with the equality condition replaced or relaxed as appropriate for the specific alignment class. Thus each alignment problem corresponds to specifying a constraint family $\mathcal{C}$ and then determining the structure of $\mathcal{E}_{\mathcal{C}}(\mathbf{t}, \mathbf{s})$.

We now describe several common tokenizers and constraint families on the function $f$. Each family defines a corresponding set of valid maps. Each example assumes a given alphabet $\Sigma$ with $S, T \in \Sigma^*$. Unless otherwise specified, the tokenizer $\tau$ is the character tokenizer, $\mathbf{t} = \tau_{\mathrm{char}}(T)$, $\mathbf{s} = \tau_{\mathrm{char}}(S)$.

**Contiguous alignment.** The strongest constraint on the alignment map $f$ is contiguity: the target must appear as a contiguous sequence in the source. The constraint family and map set are

$$\mathcal{C}_{\mathrm{CA}} = \{\, f : f(j{+}1) = f(j) + 1 \text{ for all } j \,\}, \tag{6}$$
$$\mathcal{E}_{\mathrm{CA}}(\mathbf{t}, \mathbf{s}) = \{f : f \in \mathcal{C}_{\mathrm{CA}},\ s_{f(j)} = t_j \ \forall j\}. \tag{7}$$

This class recovers exact substring search: each token of the target must match the source at consecutive positions. An example of this is classical string search [Knuth et al., 1977]. The contiguity constraint forces $f$ to be injective. In the worst case, the number of valid maps is linear in the input size when all tokens are identical ($|\mathcal{E}_{\mathrm{CA}}(\mathbf{t}, \mathbf{s})| \leq n - m + 1$).

**Ordered alignment.** Relaxing contiguity gives ordered, non-contiguous matches. As previously discussed, the constraint family and map set are

$$\mathcal{C}_{\mathrm{OA}} = \{\, f : f(1) < f(2) < \cdots < f(m) \,\}, \tag{8}$$
$$\mathcal{E}_{\mathrm{OA}}(\mathbf{t}, \mathbf{s}) = \{f : f \in \mathcal{C}_{\mathrm{OA}},\ s_{f(j)} = t_j \ \forall j\}. \tag{9}$$

This class captures ordered subsequences, allowing gaps but preserving the order of tokens. Examples include classical dynamic-programming techniques to find the longest common subsequence (LCS) [Hirschberg, 1975]. The number of alignments in the worst case is achieved when all tokens match $|\mathcal{E}_{\mathrm{OA}}(\mathbf{t}, \mathbf{s})| \leq \binom{n}{m} \approx \frac{n^m}{m!}$.

Classical global and local aligners (Needleman–Wunsch [Needleman and Wunsch, 1970], Smith–Waterman [Smith and Waterman, 1981], Wagner–Fischer [Wagner and Fischer, 1974]) operate in this same constraint family, but select a single map by maximizing a scoring function with match rewards and gap penalties instead of returning all valid maps.

$$\mathrm{Score}(f) = \sum_{j=1}^{m} S(s_{f(j)}, t_j) \; - \; \text{gap penalties}. \tag{10}$$

**Permutation alignment.** Removing monotonicity but requiring bijectivity yields exact reorderings: every source position is used exactly once (surjective) and no duplications occur (injective). The constraint family and map set are

$$\mathcal{C}_{\mathrm{PA}} = \{\, f : f \text{ bijective} \,\}, \tag{11}$$
$$\mathcal{E}_{\mathrm{PA}}(\mathbf{t}, \mathbf{s}) = \big\{ f : f \in \mathcal{C}_{\mathrm{PA}},\ s_{f(j)} = t_j \ \forall j \big\}. \tag{12}$$

This class models rearrangements such as inversions or gene-order shuffling. When $m = n$, the worst case number of valid alignments is achieved when all tokens are identical $|\mathcal{E}_{\mathrm{PA}}(\mathbf{t}, \mathbf{s})| \leq n!$.

If $f$ is injective but not surjective, then each target position selects a distinct source position, but the mapping may reorder these positions arbitrarily. This class represents subsequence alignment up to arbitrary reordering. If $f$ is surjective but not injective, then every source position is used by at least one target position, and some may be used repeatedly. Without order constraints, this class corresponds to many-to-one assignments (partitions of the target indices).

**General rearrangement alignment.** Dropping both the order constraint and cardinality constraint allows arbitrary reorderings, including duplications or deletions. Here duplication corresponds to non-injective $f$ (two targets mapping to the same source position), and deletion corresponds to non-surjective $f$ (some source positions not hit by any target). The constraint family and map set are

$$\mathcal{C}_{\mathrm{RA}} = \{\, f : f \text{ arbitrary} \,\}, \tag{13}$$
$$\mathcal{E}_{\mathrm{RA}}(\mathbf{t}, \mathbf{s}) = \big\{ f : s_{f(j)} = t_j \ \forall j \big\}. \tag{14}$$

This class captures structural processes such as large-scale genome rearrangements and other arbitrary reorderings with duplications and deletions. In the worst case, the number of alignments is achieved when every target token matches every source position ($|\mathcal{E}_{\mathrm{RA}}(\mathbf{t}, \mathbf{s})| \leq n^m$.)

If we instead use a $k$-mer tokenizer, letting $\mathbf{u} = \tau_k(S)$ and $\mathbf{v} = \tau_k(T)$, this same constraint family underlies pseudoalignment [Bray et al., 2016, Melsted et al., 2021, Sullivan et al., 2024]: a map $f$ is allowed to be arbitrary, and practical algorithms retain only membership information (whether each $k$-mer of $\mathbf{u}$ appears somewhere in $\mathbf{v}$) rather than the full map $f$.

**Approximate alignment.** Any of the classes above can be combined with a relaxed matching rule. Instead of requiring exact equality $s_{f(j)} = t_j$, we allow mismatches under a distance threshold $d(s_{f(j)}, t_j) \leq \varepsilon$ for all $j$. Here $\tau$ may be the character tokenizer or a higher-level tokenizer (e.g. words). This relaxation yields mismatch-tolerant variants of contiguous, ordered, or rearrangement alignment. Examples include Hamming-distance–based matching algorithms.

#### 1.3 A partial order of alignment

The alignment classes above differ only in the constraints they impose on the map function $f$. Stronger constraint families admit fewer maps, while weaker families admit more. This creates a natural hierarchy. Formally, for two constraint families $\mathcal{C}_1$ and $\mathcal{C}_2$, we write $\mathcal{C}_1 \preceq \mathcal{C}_2$ if any map that satisfies $\mathcal{C}_1$ also satisfies $\mathcal{C}_2$. Equivalently, $\mathcal{E}_{\mathcal{C}_1}(\mathbf{x}, \mathbf{y}) \subseteq \mathcal{E}_{\mathcal{C}_2}(\mathbf{x}, \mathbf{y})$ for all $\mathbf{x}, \mathbf{y}$.

**Theorem 1.1** (Alignment classes form a partial order)**.** *Let* $\mathsf{C}$ *be the set of alignment classes defined by constraint families. The relation* $\preceq$ *on* $\mathsf{C}$ *is a partial order: it is reflexive, antisymmetric, and transitive. Moreover, the classes introduced above satisfy* $\mathcal{C}_{\mathrm{CA}} \preceq \mathcal{C}_{\mathrm{OA}} \preceq \mathcal{C}_{\mathrm{PA}} \preceq \mathcal{C}_{\mathrm{RA}}$.

*Proof.* Reflexivity follows because every map set is contained within itself: $\mathcal{E}_\mathcal{C}(X, Y) \subseteq \mathcal{E}_\mathcal{C}(X, Y)$. For antisymmetry, if $\mathcal{C}_1 \preceq \mathcal{C}_2$ and $\mathcal{C}_2 \preceq \mathcal{C}_1$, then $\mathcal{E}_{\mathcal{C}_1}(X, Y) = \mathcal{E}_{\mathcal{C}_2}(X, Y)$, for all $X, Y$ so the two classes impose the same constraints on $f$. Transitivity is immediate from the transitivity of set inclusion.

For the chain, each step relaxes one structural condition. Contiguous alignment requires $f(j+1) = f(j) + 1$; ordered alignment keeps strict order but allows gaps; permutation alignment removes the monotonicity requirement but still forbids duplication and deletion; and the rearrangement class imposes no restrictions on $f$. Each relaxation enlarges the corresponding map set, establishing the chain. □

##### 1.4 Alignment ambiguity

Different constraint families yield map sets of different sizes. Strict classes often admit a unique map or none at all, while weaker classes may admit many. Ambiguity arises whenever the source contains repeated tokens or when positional restrictions are relaxed. Note that as a consequence of the chain more constrained mappings have fewer alignments: $|\mathcal{E}_{\text{CA}}(\mathbf{t}, \mathbf{s})| \le |\mathcal{E}_{\text{OA}}(\mathbf{t}, \mathbf{s})| \le |\mathcal{E}_{\text{PA}}(\mathbf{t}, \mathbf{s})| \le |\mathcal{E}_{\text{RA}}(\mathbf{t}, \mathbf{s})|$.

For example, for $s = \texttt{DAD}$ and $t = \texttt{ADD}$, the positions matching each `D` in $t$ may map to either occurrence of `D` in $s$. Under ordered alignment, no map exists, but under permutation or rearrangement constraints there are multiple valid choices for $f$. Thus relaxing constraints not only enlarges the map set but also increases the number of admissible mappings; a larger map set corresponds to greater alignment ambiguity.

#### 2 Discussion

Historically, alignment methods in computational linguistics and genomics focused on developing algorithms for a specific task. These algorithms were optimized independently for biological sequences or natural language text. As a result, similarities between these algorithms remained silo-ed in different journals and software tools.

We unify these approaches by framing alignment as a map $f$ between token sequences, with methods primarily distinguished by the constraints imposed on this map. Under this view, any alignment method is defined by three choices: a tokenizer, a constraint family on the map, and a selection rule for choosing an alignment.

This framing makes relationships between methods explicit. For example, classical text alignment imposes stricter constraints when mapping an extracted piece of text to a passage than genomics pseudoalignment does when mapping a sequencing read to a transcriptome. These constraints implicitly define a trade-off between computational complexity and alignment ambiguity. Once an algorithm is placed within a constraint family, its possible relaxations or tightenings become clear, along with their effects on ambiguity and computational cost. This allows scientists to reason about speed, memory usage, and alignment multiplicity prior to implementation.

By organizing alignment methods in this way, we connect two disciplines that have largely evolved independently. This connection enables mathematical and algorithmic ideas developed in genomics to be reused in natural language processing, and vice versa (Table 1). Indexing strategies, hashing schemes, and alignment selection rules can be transferred across domains within a shared formal framework. In this work, we combine algorithms and computational techniques from genomics [Bray et al., 2016, Melsted et al., 2021, Sullivan et al., 2024] with tokenization and alignment selection ideas from natural language processing [Hirschberg, 1975, Sennrich et al., 2016]. The alignment tool discussed in the manuscript (`taln`) emerges naturally from this framework. It uses ordered alignment over subword tokens to balance alignment flexibility with computational efficiency.

### 3 Simple alignment examples

We list short examples that illustrate how different constraint families $\mathcal{C}$ behave on small strings. For a source $\mathbf{x}$ and target $\mathbf{y}$, we describe whether maps exist under each class and, when helpful, the number of valid maps.

**Contiguous alignment.** Requires $f(j+1) = f(j) + 1$.

- $\mathbf{x} = \texttt{SCATTER}$, $\mathbf{y} = \texttt{CAT}$: one valid map $\{(2, 3, 4)\}$.
- $\mathbf{x} = \texttt{ABCDE}$, $\mathbf{y} = \texttt{ACE}$: no contiguous map.

**Ordered alignment.** Requires $f(1) < \cdots < f(m)$.

- $\mathbf{x} = \texttt{ABCDE}$, $\mathbf{y} = \texttt{ACE}$: one map $\{(1, 3, 5)\}$, which is ordered but non-contiguous.
- $\mathbf{x} = \texttt{DAD}$, $\mathbf{y} = \texttt{ADD}$: no map.

**Permutation alignment (bijective)** Here $f$ is bijective and order is unrestricted.

- $\mathbf{x} = \texttt{ABC}$, $\mathbf{y} = \texttt{CBA}$: one map, the reversal $\{(3, 2, 1)\}$.
- $\mathbf{x} = \texttt{ABA}$, $\mathbf{y} = \texttt{BAA}$: two maps $\{(2, 1, 3), (2, 3, 1)\}$.

Relaxing bijectivity yields closely related classes:

- *Injective but not surjective* (unordered subsequence alignment).
- *Surjective but not injective* (sequence partition).

**General rearrangement alignment.** Here $f$ is arbitrary (not required to be bijective nor injective).

- $\mathbf{x} = \texttt{CAT}$, $\mathbf{y} = \texttt{CAAT}$: duplication $(1, 2, 2, 3)$, since the single `A` in $\mathbf{x}$ must be used twice.
- $\mathbf{x} = \texttt{GGA}$, $\mathbf{y} = \texttt{GA}$: deletion $\{(1, 3), (2, 3)\}$, since one of the `G` positions in $\mathbf{x}$ is not used.

With a $k$-mer tokenizer, this rearrangement class underlies pseudoalignment. With $k$=2,

- $\mathbf{x} = \texttt{GATTACA}$ yields 2-mers $\mathbf{u} = \texttt{GA,AT,TT,TA,AC,CA}$,
- $\mathbf{y} = \texttt{TTA}$ yields 2-mers $\mathbf{v} = \texttt{TT,TA}$.

An arbitrary map $f$ with $u_{f(j)} = v_j$ exists (both `TT` and `TA` occur in $\mathbf{u}$), so $\mathbf{y}$ is pseudoaligned to $\mathbf{x}$ with the alignment $\{(3, 4)\}$.

**Table 1:** Alignment classes defined by structural constraints on the map function $f$.

| **Name** | **Constraint family** $\mathcal{C}$ | **Map set** $\mathcal{E}_{\mathcal{C}}(\mathbf{t},\mathbf{s})$ | **Classical algorithms** |
|---|---|---|---|
| **Contiguous alignment** | $\{f : f(j{+}1) = f(j) + 1\}$ | $\{f : f \in \mathcal{C}_{\mathrm{CA}},\ s_{f(j)} = t_j\ \forall j\}$ | KMP, Boyer–Moore; exact substring search |
| **Ordered alignment** | $\{f : f(1) < \cdots < f(m)\}$ | $\{f : f \in \mathcal{C}_{\mathrm{OA}},\ s_{f(j)} = t_j\ \forall j\}$ | LCS, edit distance, Needleman–Wunsch, Smith–Waterman |
| **Permutation alignment (bijective)** | $\{f : f \text{ bijective}\}$ | $\{f : f \in \mathcal{C}_{\mathrm{PA}},\ s_{f(j)} = t_j\ \forall j\}$ | Genome inversion/transposition models; permutation distance |
| **General rearrangement** | $\{f : f \text{ arbitrary}\}$ | $\{f : s_{f(j)} = t_j\ \forall j\}$ | Genome rearrangements, V(D)J recombination; pseudoalignment (with $k$-mers) |
